# Effects of rising amyloidβ levels on hippocampal synaptic transmission, microglial response and cognition in *APP*_Swe_/*PSEN1*_M146V_ transgenic mice

**DOI:** 10.1101/420349

**Authors:** Evelyn Medawar, Tiffanie Benway, Wenfei Liu, Taylor A. Hanan, Peter Haslehurst, Owain T. James, Kenrick Yap, Laurenz Muessig, Fabia Moroni, Muzzamil A. Nahaboo Solim, Gaukhar Baidildinova, Rui Wang, Jill C. Richardson, Francesca Cacucci, Dervis A. Salih, Damian M. Cummings, Frances A. Edwards

**Affiliations:** University College London, Gower Street, London, WC1E 6BT, UK; Neurosciences Therapeutic Area, GlaxoSmithKline R&D, Gunnels Wood Road, Stevenage, SG1 2NY, UK

**Keywords:** Alzheimer’s disease, Dementia, mouse model, synaptic transmission, microglia, plaque, neurodegeneration

## Abstract

**Background:** Progression of Alzheimer’s disease is thought initially to depend on rising amyloidβ and its synaptic interactions. Transgenic mice (TASTPM; *APP*_Swe_/*PSEN1*_M146V_) show altered synaptic transmission, compatible with increased physiological function of amyloidβ, before plaques are detected. Recently, the importance of microglia has become apparent in the human disease. Similarly, TASTPM show a close association of plaque load with upregulated microglial genes.

**Methods:** CA1 Synaptic transmission and plasticity were investigated using *in vitro* electrophysiology. Migroglial relationship to plaques was examined with immunohistochemistry. Behaviour was assessed with a forced-alternation T-maze, open field, light/dark box and elevated plus maze.

**Findings:** The most striking finding is the increase in microglial numbers in TASTPM, which, like synaptic changes, begins before plaques are detected. Further increases and a reactive phenotype occur later, concurrent with development of larger plaques. Long-term potentiation is initially enhanced at pre-plaque stages but decrements with the initial appearance of plaques. Finally, despite altered plasticity, TASTPM have little cognitive deficit, even with a heavy plaque load, although they show altered non-cognitive behaviours.

**Interpretation:** The pre-plaque synaptic changes and microglial proliferation are presumably related to low, non-toxic amyloidβ levels in the general neuropil and not directly associated with plaques. However, as plaques grow, microglia proliferate further, clustering around plaques and becoming phagocytic. Like in humans, even when plaque load is heavy, without development of neurofibrillary tangles and neurodegeneration, these alterations do not result in cognitive deficits. Behaviours are seen that could be consistent with pre-diagnosis changes in the human condition.

**Research in context:** *Evidence before this study:* There is a large body of research examining many aspects of phenotypes associated with mouse models of Alzheimer’s disease – a PubMed search for the terms Alzheimer* AND mouse returns in excess of 21000 articles. However, there are few systematic articles pulling together pathological, functional (electrophysiological), and behavioural analyses across the life-span of such models. There is also a number of conflicting outcomes, for example reports of impaired versus enhanced synaptic plasticity; cognitive impairments or not. Recently, the importance of microglia in Alzheimer’s disease has come to the fore in human Genome Wide Association Studies (GWAS), with variants of a number of microglial genes identified as risk-factors for developing the disease. Interestingly, we have recently reported that *Trem2* and other genes identified as risk-factors in humans are strongly up regulated in close association to plaque development in the mouse model used in this study. Moreover, this previous study predicted two of the most recently identified genes that were identified in GWAS since the publication of our paper. We have previously used this model to identify the earliest synaptic changes and shown changes in release of glutamate, the primary excitatory neurotransmitter in the brain, to occur even before plaques are detectable.

*Added value of this study:* By studying this transgenic mouse model of Alzheimer’s disease, throughout the development of plaques, from prior to detection through to heavy plaque loads, we have been able to identify a clear time course of key phenotypic changes associated with early disease. In particular, this study identifies the very early changes in microglia and can separate the time course of the microglial phenotype. In addition, we detail the changes in synaptic plasticity over time and importantly identify that, like in humans in the absence of Tau tangles or neurodegeneration, considerable synaptic changes can occur and a heavy plaque load without resulting in substantial cognitive loss.

*Implications of all the available evidence:* Our data indicate that rising amyloid beta prior to detectable plaque deposition results in changes in synaptic function that likely reflects an enhanced physiological effect of amyloid beta. At this stage, microglia proliferate but do not activate. Once plaques begin to appear, microglia migrate to surround the plaque and become phagocytic, likely targeting dystrophic synapses and neurites caused by the cloud of highly-toxic amyloid beta around the plaque. Similarly to humans, who have plaques but no tangles and have yet to develop substantial neurodegeneration, cognitive deficits are not seen, even with a heavy plaque load; behavioural changes are limited to anxiety-like effects. This investigation of the parallel time-course of events highlights the probability that, if progression of disease can be reversed or slowed early enough, before Tau tangles and substantial neurodegeneration occur, the symptoms of cognitive decline could be very largely avoided. Moreover, it suggests that the substantial increases in microglia number and upregulation of their specific gene expression in association with plaques, is not associated with cognitive loss and may indeed be protective.

## Introduction

The onset and progression of Alzheimer’s disease (AD) is most likely initiated by environmental factors interacting with predisposed genetic risks, many of which have now been identified in genome-wide association studies (see reference 1 for review). In familial AD, an inherited mutation causes rising amyloidβ (Aβ) levels and it has long been established that this triggers a chain of events that leads to the eventual cognitive decline.^[2]^ In sporadic AD, it is highly likely that initial triggering events (genetic and/or environmental) also lead to rising Aβ and that, like in the familial disease, Aβ levels represent an essential contributor to the ongoing pathology and eventual neurodegeneration.^[3, 4]^ Under normal physiological conditions, Aβ acts as an activity-dependent synaptic modulator which, when released from presynaptic neuronal terminals, increases probability of glutamate release.^[5]^ Inappropriate neuronal activity and/or genetic imbalances in production versus clearance mechanisms may thus lead to a prolonged rise in Aβ levels,^[6]^ enabling the formation of Aβ oligomers and deposition of plaques. It is notable that restoring γ-frequency oscillations in transgenic mouse models expressing genes harbouring familial mutations associated with familial AD, reduces Aβ load^[7]^ and restores cognitive deficits.^[8]^

Animal models for AD have generally depended on transgenic expression, or more recently knock-in of the gene variants that cause the familial dominantly inherited forms of the disease, particularly amyloid precursor protein (*APP*) or presenilin 1 or 2 (*PSEN1* or *PSEN2*). This is an effective approach for initiating the rise in Aβ and the deposition of plaques, albeit by a different trigger than in sporadic AD. According to the ‘amyloid hypothesis’, in the human disease, rise in Aβ and plaque deposition is suggested to lead to altered neuronal ionic homeostasis and increased oxidative stress. Together, these may result in increased kinase activity on microtubule-associated protein Tau, resulting in Tau hyperphosphorylation and neurofibrillary tangle formation.^[9]^ The exact mechanisms linking Aβ and Tau phosphorylation remain unclear and unfortunately, despite the initiation of Aβ pathology, mouse models with genes for familial Alzheimer’s disease do not completely recapitulate these later events. As documented in the initial descriptions of the TASTPM mice, which are hemizygous for both human APP_Swe_ and human PSEN1_M146V_, plaques are first detected at approximately 4 months of age and a considerable plaque load develops by 8 months.^[10, 11]^ Furthermore, phosphorylation of Tau is detectable in the dystrophic neurites around plaques,^[11]^ recently suggested to be an early stage of Tau pathology.^[12]^ An inability to extinguish hippocampal-dependent contextual fear conditioning at 4 months when plaques are first detected^[13, 14]^ and a deficit in novel object recognition at 6 months,^[15]^ have also been reported.

We recently reported alterations in synaptic transmission preceding the detection of plaques, manifesting as a loss of spontaneous action potentials in Schaffer collateral axons and a concomitant increase in probability of glutamate release.^[16]^ Importantly, we have demonstrated an almost 1:1 correlation of plaque load with expression of a module of microglial genes throughout the life of these and other transgenic mice. This correlation in Αβ mice contrasts with the interaction of microglia and neurofibrillary tangles in mice with Tau mutations, in which microglial genes are only upregulated with advanced tangle load (www.mouseac.org).^[10]^

Here we extend our previous studies on TASTPM mice to understand the relationship between early changes in synaptic transmission, synaptic plasticity, cognitive function and microglia. We study the development of plaques in more detail and dissect out the microglial response to distinguish between the numbers of microglia and their phagocytic status. We find that, like synaptic changes, microglia are more prevalent even before plaques are detectable, whereas their phagocytic phenotype is age-related, coming much later. We then proceed to study hippocampal synaptic plasticity and hippocampus-dependent learning, in the forced-alternation T-maze, enabling identification of Aβ phenotypes, finding little cognitive deficit even with a heavy plaque load but clear behavioural changes, probably related to increased levels of anxiety. While we continue to focus mainly on the previously defined TASTPM mice (double hemizygote), we have broadened the study to investigate dose dependency of the transgene by including mice homozygous for both genes. The effect of the individual genes is also investigated in mice with hemizygous expression of only one or the other gene. When not otherwise defined, TASTPM mice hence refers to the double hemizygous mouse.

## Methods

### Animals

All experiments were performed in agreement with the UK Animals (Scientific Procedures) Act 1986, with local ethical approval and in agreement with the GlaxoSmithKline statement on use of animals. Male TASTPM mice and C57Bl/6j mice were supplied by GlaxoSmithKline and bred either at Charles River Laboratories International, Inc. (Margate, UK) or at UCL by crossing male homozygous TASTPM with female C57Bl/6j. Age-matched, non-littermate male C57Bl/6j mice were used as wild type controls. In some experiments, double homozygous TASTPM were bred. Single mutant TAS (APP_Swe_)^[17]^ or TPM (PSEN1_M146V_)^[15]^ mice were bred by crossing hemizygous parents. Mice from Charles River were shipped to UCL upon weaning at 21-days-old.

In this study we avoided single housing by keeping mice in large open cages (20 × 35 × 45 cm) and enriching their environments. Under these conditions, while the aggressive nature of the TASTPM mouse is not completely avoided, it is less of a problem and allows group housing to be maintained over the lifetime of the mice. Thus, cages containing 2-8 male mice were maintained in a 12-hour light/12-hour dark cycle with food (Envigo 2018 Teklad global 18% protein rodent diet) and water *ad libitum*. Environmental enrichment consisted of changes of food location, bedding type (e.g. tissue, shredded paper, paper roll, paper bags) and inanimate objects (e.g. running wheels, rodent balls, tubing, houses (mostly purchased from Eli Lilly Holdings Limited, Basingstoke, UK)) within the cage at least once per week. Mice were used for experimentation at the ages stated (± 0·5 months) and, where unavoidable, were single-housed for no longer than 24 hours. Tails or ear punches were used for genotyping by standard PCR protocols to ensure the presence of the expected genes.

## Genotyping

### Genotype confirmation using conventional PCR methods

Briefly, genomic DNA was extracted using the ‘HotSHOT’ lysis method. Alkaline lysis reagent (25 mM NaOH, 0·2 mM EDTA, pH12) was added to tissue samples prior to heating to 95°C for 30 minutes. The sample was then cooled to 4°C before the addition of neutralisation buffer (40 mM Tris-HCl, pH 5). The PCR reaction was performed through addition of MyTaq DNA Polymerase (Bioline) reaction buffer and primer pairs:

TAS (APP_Swe_):

5’ GAATTGACAAGTTCCGAGGG 3’
5’ GGGTACTGGCTGCTGTTGTAG 3’ TPM (PSEN1_M146V_):

5’ GTTACCTGCACCGTTGTCCT 3’
5’ GCTCCTGCCGTTCTCTATTG 3’

using the cycling parameters: 94°C (30 s), 58°C (30 s), 72°C (30 s), for 30 cycles and a final extension at 72°C for 4 min. PCR product sizes 366 bp for TAS and 104 bp for TPM.

## Immunohistochemistry

Animals were deeply anaesthetised (1:10 Euthatal:Intra-Epicaine, National Veterinary Supplies) and transcardially perfused with 0·1 M phosphate buffer saline (PBS) followed by 10% buffered formal saline (Pioneer Research Chemicals Ltd). Alternatively, single hemispheres were drop-fixed immediately following brain extraction for electrophysiology. The brains were post-fixed in 10% buffered formal saline for 24hrs and cryoprotected in 30% sucrose/0·03% sodium azide/PBS at 4°C for at least 24hrs before sectioning or storage. Transverse sections were cut at 30 μm through the full left hippocampus using a frozen sledge microtome (SM 2000 R, Leica) and collected into a 24-well plate containing PBS/sodium azide (0·03%) for storage at 4°C. Serial sections were placed in separate wells until all wells contained a section and collection then continued serially from Well 1 so that within each well the transverse sections were from the length of the hippocampus at least 720μm apart.

Standard immunohistochemical techniques were employed. For Aβ staining only, antigen retrieval was achieved by submerging sections in 10 mM sodium citrate (pH 6·0) in 0·05% Triton X-100 and heated in a water bath at 80°C for 30 minutes. Sections for all immunohistochistry were then washed in PBS, followed by 0·3% Triton X-100 in PBS and subsequent blocking in 8% horse serum/Triton/PBS for 1 hour. Incubation with primary antibody (table 1) in blocking solution was performed overnight at 4°C. Sections were again washed with Triton/PBS. The appropriate Alexa-conjugated secondary antibody (1:500; Invitrogen) was added to blocking solution for a 2-hour incubation at room temperature in the dark. Following PBS wash, DAPI (1:10,000) was applied to all sections for 5 minutes. Sections were washed for a final time in PBS before mounting. Age-matched sections from wild type controls were stained in parallel for all ages. Sections were mounted in anatomical order onto SuperFrost Plus glass slides by floating on PBS and then cover-slipped using Fluoromount G mounting medium.

**Table 1:** *Antibodies used for immunohistochemistry:* Ms Evelyn Medawar MSc*

| Antibody | Dilution | Source | RRID |
| --- | --- | --- | --- |
| IBA1 | 1:500 | Wako (019-19741) | AB_839504 |
| CD68 | 1:500 | BioRad (MCA1957) | AB_322219 |
| Aβ1-40 | 1:300 | ThermoFisher (44-136) | AB_2533599 |

### Imaging and data analysis

Sections were imaged for quantification using an EVOS FL Auto Cell Imaging System (Life technologies). Tiled images were taken of the whole transverse hippocampal section using a 20X objective. To determine cell densities, an area of 400 μm × 240 μm was defined in the CA1, CA3 and the inner blade of the dentate gyrus. Cell counts were performed using Adobe Photoshop CS6. A minimum of three sections were used to create a mean for each animal. Sections for any given condition were obtained from the same collection well within the 24-well plate and were therefore a minimum of 720 μm apart, thus avoiding multiple counts of the same cells. Counts of objects touching the boundaries of the area of interests were only included from the north and east borders and excluded from the south and west borders.

## Electrophysiological recordings

### Acute hippocampal brain slice preparation

Mice were decapitated and the brain rapidly removed and placed in ice-cold dissection artificial cerebrospinal fluid (ACSF, containing (in mM): 125 NaCl, 2·4 KCl, 26 NaHCO_3_, 1·4 NaH_2_PO_4_, 20 D-glucose, 3 MgCl_2_, 0.5 CaCl_2_, pH 7·4, ~315 mOsm/l). After approximately two minutes in ice-cold dissection ACSF, the brain was prepared for slicing by removing the cerebellum, hemisection of the forebrain and a segment cut away from the dorsal aspect of each hemisphere at an angle of approximately 110° from the midline surface to optimise slicing transverse to the hippocampus. Each hemisphere was then glued with cyanoacrylate (Loctite 406, Henkel Loctite Limited, UK) on this surface onto the stage of a vibrating microtome (Integraslice model 7550 MM, Campden Instruments, Loughborough, UK) containing frozen dissection ACSF and 400 μm transverse slices of hippocampus cut. As each slice of a hemisphere was cut, the hippocampus was dissected out, retaining a portion of entorhinal cortex and the resulting smaller slice was placed into a chamber containing ‘Carbogenated’ (95% O_2_/5% CO_2_; BOC Limited) dissection ACSF at room temperature (approximately 21°C). After 5 minutes, slices were then transferred into a fresh chamber held at 36°C with the same dissection ACSF. At 5-minute intervals, they were then consecutively transferred to physiological Ca^2+^ and Mg^2+^ ion concentrations (in mM): i) 1 Mg^2+^, 0.5 Ca^2+^; ii) 1 Mg^2+^, 1 Ca^2+^; iii) 1Mg^2+^, 2 Ca^2+^. After approximately 20 minutes at 35°C (i.e., once transferred into the 1 Mg^2+^, 2 Ca^2+^ ACSF).

### Patch-clamp recordings in brain slices

Once transferred to 1 Mg^2+^, 2 Ca^2+^ ACSF, slices were allowed to return to room temperature and after at least a further 40 minutes recovery time, a single slice was transferred to a submerged chamber and superfused with recording ACSF (containing (in mM): 125 NaCl, 2·4 KCl, 26 NaHCO_3_, 1·4 NaH_2_PO_4_, 20 D-glucose, 1 MgCl_2_, 2 CaCl_2_, bubbled with Carbogen). Individual CA1 pyramidal or dentate gyrus granule neurones were visualised using infrared-differential interference contrast microscopy on an upright microscope (model BX50WI, Olympus, UK). Glass microelectrodes for patch-clamp were pulled from borosilicate glass capillaries (Catalogue number GC150F-7.5, 1·5 mm OD × 0·86 mm ID, Biochrom-Harvard Apparatus Ltd, Cambridge, UK) on a vertical puller (model PP830, Narishige International Ltd, London UK). Electrodes (tip resistance approximately 5 MΩ) were filled with a CsCl-based internal solution (containing (in mM): CsCl 140, HEPES 5, EGTA 10, Mg-ATP 2, pH 7·4, ~290 mOsm/l). Patch-clamp recordings were performed using an Axopatch 1D (Molecular Devices, Sunyvale, CA, USA), and current signals low-pass filtered at 10 kHz then 2 kHz (Brownlee Precision Model 440, NeuroPhase, Santa Clara, CA, USA) during digitization (10 kHz; 1401plus, Cambridge Electronic Design, Limited, Cambridge, UK) and acquired using WinWCP (for isolated events; version 4.6.1; John Dempster, Strathclyde University, UK) and WinEDR (for continuous recordings; John Dempster, Strathclyde University, UK). Stimulation was applied *via* a patch electrode filled with ACSF, placed extracellularly in the appropriate axon path using a square pulse constant-voltage stimulator (100 μs; DS2A-MkII, Digitimer Ltd, UK) triggered by WinWCP.

WinEDR synaptic analysis software was used for detection of spontaneous and miniature currents and WinWCP used to analyse identified spontaneous, miniature and evoked currents. Criteria for detection of spontaneous or miniature currents was to remain over a threshold of 3 pA for 2 ms. Currents were inspected by eye and only included if the rise time was <3 ms and faster than the decay.

### Field potential recordings in brain slices

Slices were transferred as needed to a heated (30±1°C) submerged chamber and superfused with ACSF and allowed to recover for 1 h in the recording chamber. A glass stimulating electrode (filled with ACSF, resistance ~2 MΩ) and an identical recording electrode (connected to an AxoClamp 1B via a 1X gain headstage) were both positioned in stratum radiatum of the CA1 field to obtain a dendritic excitatory postsynaptic field potential (fEPSP). Recordings were controlled and recorded using WinWCP software (as above), filtered at 10 kHz and subsequently at 3 kHz and digitized at 10 kHz via a micro1401 interface (Cambridge Electrical Designs, UK). Stimuli (constant voltage 10-70V, 100 μs; model Digitimer DS2A-MkII or Grass SD9) were applied at 0·1 Hz and resultant fEPSPs subsequently averaged over consecutive 1-minute intervals. Stimulation intensity was set at approximately 30-50% of the intensity required to evoke a population spike or the maximum fEPSP amplitude obtained and a ≥15-minute stable baseline recorded. LTP conditioning was applied at test-pulse stimulus intensity and consisted of either 3 trains of tetani, each consisting of 20 pulses at 100 Hz, 1·5 s inter-train interval or 4 trains of theta-burst stimuli (TBS), each train consisting of 4 pulses at 100 Hz repeated 8 times at 20 Hz; inter-train interval 1 minute. Following conditioning, fEPSPs were evoked at 0·1 Hz for 1 hour.

## Behavioural testing

### T-maze forced alternation task

Previously reported methods, optimised for mouse, were used to assess hippocampus-dependent learning ^[18]^. Mice were food deprived to 90% free-feeding-weight, beginning 2 days before the start of the habituation phase and with *ad libitum* access to water. Each mouse was handled at the start of food deprivation and throughout T-maze habituation for 15-20 minutes per weekday.

The T-maze was constructed from three arms, each measuring 50 × 8 cm with 10 cm colourless Perspex walls and a grey floor, mounted on a table in the centre of a room with numerous distal visual cues, such as black and white posters on the walls. Black barriers were used to block the start and goal arms. Reward consisted of a drop of Nestlé Carnation^TM^ condensed milk that was placed at the end of each goal arm. Arms were cleaned with 70% ethanol between all runs to reduce odour cues. In addition, in an inaccessible well a drop of reward is always present in both arms.

Mice received 4 days of habituation to the maze, during which time they were allowed to explore the maze for 5 minutes with all arms open. During the first two days of habituation, reward was scattered along the floor and in food wells to encourage exploratory behaviour; then restricted to only the food wells at the ends of the goal arms in the last two days.

The behavioural regime lasted for three weeks, with five days of training per week. Each day, animals received six trials; each trial consisted of a sample and choice run. In the sample run, one arm was blocked off. The mouse was placed at the starting point at the base of the T, the barrier was removed and the mouse was allowed to go to the available arm and given 20 s to eat a drop of reward from the food well. For the choice run, the mouse was immediately returned to the starting point and the barrier in the previously blocked arm removed. The starting barrier was then raised and the animal allowed to choose between the two arms but only rewarded if it chose the previously unvisited arm. Thus, a correct choice was scored when the mouse selected the arm not visited in the sample run. After the choice run, the mouse was removed from the maze and placed in its holding box. The location of the sample arm (left or right) was varied pseudorandomly across the session and mice received three left and three right presentations, with no more than two consecutive trials with the same sample location. Animals were allowed a maximum of 5 minutes to make a choice to enter a goal arm in both runs before a trial was aborted. If an incorrect arm was chosen during the choice run, the mouse was confined in the arm with no reward for 20 s and then removed from the maze.

During the first two weeks of training the choice run followed immediately after the sample (test) run (there was a delay of approximately 15 s between runs for cleaning and resetting the maze). Data was analysed in blocks of 2 days and hence blocks 1-5 represent the first 2 weeks of training.

On the first day of the third week, mice received a repeat of the previous training sessions in order to assess retention of the task (block 6). On the following four days, longer delays (2-10 minutes) were introduced between the sample and choice runs to extend the time that the previous choice was to be held in memory (blocks 7 and 8 in response times). During these intervals, each animal was placed in a separate holding box. Each mouse received two of each of the delay periods per day, varied pseudorandomly both within and across days. Response times were calculated from the time that the starting block was removed until the mouse made a choice of arms and all four paws had crossed the entry point. Squads of 15-17 mice were run per day. During training data are presented as blocks averaged across 2 days for each animal and expressed as mean ± SEM. For delays the four runs for each delay are averaged.

### Elevated plus maze

The plus-maze was constructed from two enclosed arms (30 cm × 5 cm × 20 cm) and two open arms (30 cm × 5 cm × 0.8 cm) connected by a small central platform (5 × 6 cm); the two closed and two open arms were positioned opposite each other, respectively. The maze was elevated 30 cm above a table.

Mice were placed in the centre of the plus maze facing an open arm and allowed to freely explore the maze for one 6-minute trial. The time spent on each arm (open *versus* closed) was recorded, as well as the number of entries into the arms. An entry into an arm was defined as all four paws resting on a given arm.

### Open field

The open field consisted of a plastic cylinder (diameter: 47·5 cm, height 36 cm) with a white plastic floor. Mice were placed on the periphery of the open field floor and allowed to explore freely for 30 minutes. The path of each mouse was recorded using dacQUSB recording system (Axona, St. Albans, U.K) at a sample rate of 50 Hz. The open field was optically divided into a central circle and a peripheral ring, each with equal area. The path of the mouse was analysed offline using ImageProPlus. Path length and dwell times in the periphery and centre were calculated using custom made routines written in Matlab R2010a (MathWorks).

### Light/dark box

The dark box^*[19]*^ measured 20 cm × 20 cm × 30 cm, with black walls, floor and lid. The light box measured 30 cm × 30 cm × 30 cm with a white floor and light grey walls. The boxes were connected by an opening in the partition between the two compartments. An overhead light provided bright illumination in the light box. Mice were placed in the centre of the light compartment facing away from the opening and then allowed to explore for a period of 6 minutes. Time spent in each box and the number of entries into each box were recorded. An entry into a box was defined as all four paws resting inside the given box.

## Statistics

All data analysis was carried out blind to genotype. All statistics were performed using Graphpad Prism 6 with appropriately designed two-tailed t-test or ANOVA. Post hoc tests were only performed if a significant interaction between the independent variables was obtained. Animals were considered as independent samples and, where multiple data were collected from an animal, these were averaged (mean) prior to pooling. Thus sample sizes represent the number of animals. Unless stated otherwise, data are presented as mean ± SEM and differences considered significant at p<0·05.

## Results

### Plaque development in hippocampus

In TASTPM mice of different ages, individual plaques were counted in the hippocampus and their sizes measured to assess whether the distribution changed over time (Fig 1). Initially, at 3-4 months of age, only small plaques could be detected (<100 μm^2^) and these were very sparse (Fig. 1a-c). However, the number of small plaques increased exponentially, initially approximately doubling every month, and continuing to increase at a slower rate until 14 months of age (Fig. 1b). Hence, at least up until this age, new plaques were being seeded. By 7 months plaques up to 200 μm^2^ were consistently seen but were similar in frequency to the smallest plaques at 3-4 months, presumably representing the growth of the earliest seeds. Plaques of >500 μm^2^ were only consistently detected at ages over 10 months but, from this age on, a wide range of plaques sizes were evident, with occasional plaques of over 2500 μm^2^ detected. Interestingly, between 14 and 15 months of age, very little change in the number of plaques was detected but the largest plaques continued to grow. As individual plaques were not tracked over time, we cannot be entirely sure that the progression up to 14 months is due to continuous seeding of small plaques that gradually grow with ongoing plaque deposition; however, this seems the most likely explanation. The fact that the rate of addition of small plaques decreases gradually until it plateaus around 14 months of age suggests that, as the plaques get denser, some of the small plaques start to fuse with larger plaques. In some cases, large plaques were present that possessed a halo of Aβ around the dense core, with dense puncta of Aβ visible within the halo, possibly representing more recently seeded plaques being engulfed as the large plaque grows (Fig 1a).

**Figure 1.**
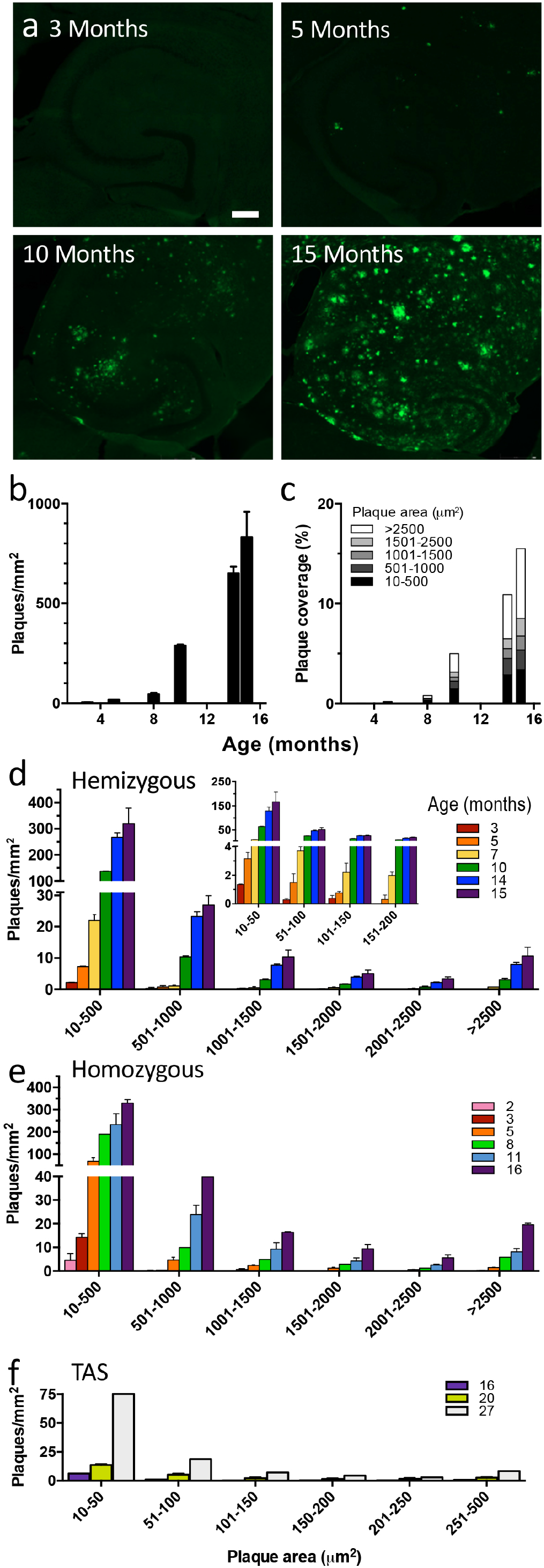
Plaque development in TASTPM mice. a) Representative micrographs of Aβ40 (green) immunofluorescence in hippocampus from 3- to 15-month-old hemizygous TASTPM mice. Scale bar 250 μm. b) Plaque density measured at different ages in TASTPM mice (n=2-3 animals per age group). c) Proportion of plaques of indicated sizes contributing to increased percentage coverage of the hippocampus with increased age. d) Density of plaques within specific size ranges (as indicated on x-axes) over age (colours as indicated in key). The inset covers smaller bin sizes for plaques up to 200 μm^2^. Note the split y-axes. e) Density of homozygous TASTPM plaques within specific size ranges (as indicated on x-axes) over age (colours as indicated in key). f) Density of TAS plaques within specific size ranges (as indicated on x-axes) over age (colours as indicated in key). Note the different x and y-axes in f compared to d and e, with few plaques and maximum plaque area under 500 μm^2^. All data are represented by mean±SEM.

When the gene-dose was doubled by using double homozygous TASTPM mice, plaques were first detected at about 2 months of age (Fig 1e). Similarly to the hemizygous mice, plaque density and size increased with age but perhaps surprisingly, did not reach a greater density than that observed in the hemizygotes.

Mice carrying only the single mutations were also examined. In TAS mice (APP_Swe_), small plaques were detected at 16 months. While plaque density and size increased with age, plaque loads did not reach that seen in the double mutant TASTPM mice, even at the very advanced age of 27 months. In TPM mice (PSEN1_M146V_), there were no plaques observed at any age.

### Microglia

Having previously observed that a module of microglial genes increased in expression in Αβ mice (including TASTPM) in close correlation with plaque load^[10]^, we went on to examine the densities of total and CD68 positive of microglia in more detail in these mice (Fig 2). There was an increased density of microglia (assessed by Iba1 positive cells) in CA1 at very early ages, with increases (approximately double) compared to wild type already evident by 2 months of age, even before plaques could be detected (Fig 2a&c; 2-8 months, n=6-12, two-way ANOVA, p<0·01). However, note that Aβ levels are already raised by 2 months^[16]^ and it is possible that small plaque seeds (< the 10 μm^2^ threshold) may already be depositing at this time. There was also an age-dependent increase in Iba1 positive microglia in both wild type and TASTPM mice at around 10 months of age.

**Figure 2.**
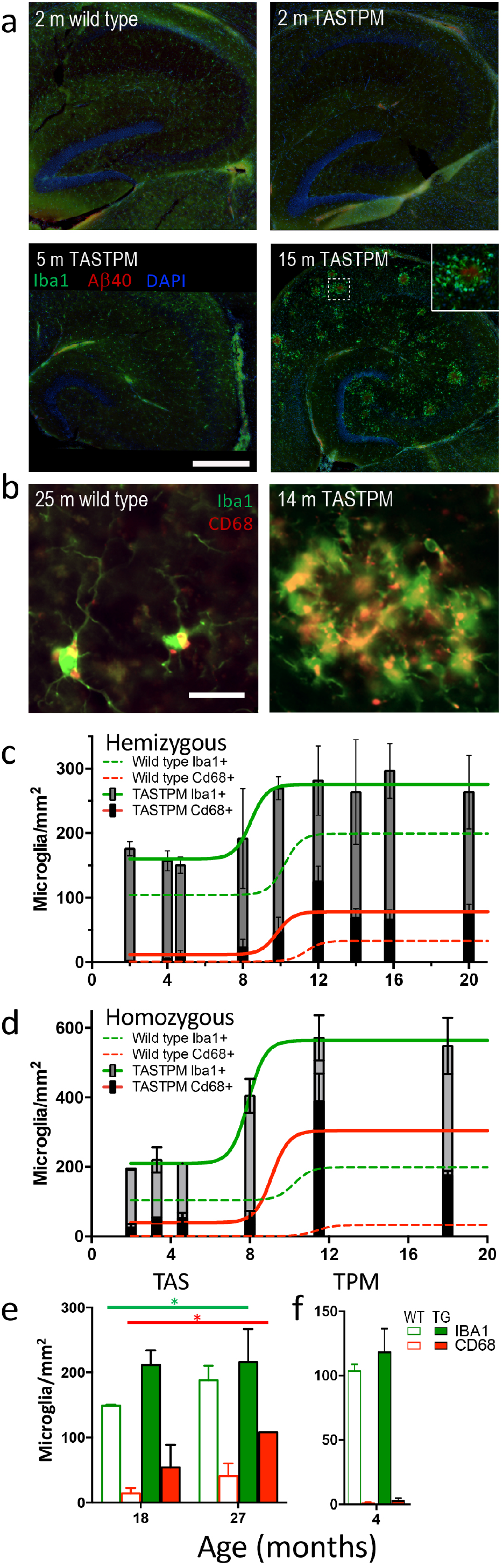
Microglial proliferation and activation in TASTPM mice. a) Microglia (Iba1, green) cluster around plaques (Aβ1-40, red) in TASTPM mouse hippocampus (nuclei: DAPI, blue). Scale bar 250 μm. b) Examples of phagocytic microglia (CD68, red + Iba1, green; yellow when colocalised) in wild type and TASTPM hippocampus. Scale bar 25 μm. c) Total microglial density (Iba1+, full bar height) and phagocytic microglial (Iba1 + CD68+, black bar) density in hemizygous TASTPM mice with age. Sigmoidal fits are shown (Iba1 green; CD68 red): unbroken lines, TASTPM; dashed lines, wild type (only fit shown for wild type). d) Total and phagocytic microglial densities in homozygous TASTPM mice with age. Sigmoidal fits are shown. e) Total and phagocytic microglial densities in the APP_Swe_ TAS mice at the ages shown. Significant main effects of genotype for both IBA1 and CD68 are indicated by *p<0.05. f) Densities of total microglia and CD68 positive phagocytic microglia in 4 month old PSEN1_M146V_ TPM microglia. All data are represented by mean±SEM.

In contrast to total microglia, the microglial phagocytic phenotype, as measured by CD68 labelling, was very low in all young animals and only increased from around 10-12 months. Again, this happened in both wild type and TASTPM mice. The percentage of phagocytic microglia tended to be slightly higher in TASTPM mice as compared to wild type (14-20 months, n=5-9, two-way ANOVA p=0·07). This coincided with the appearance of large plaques (Fig 1c&d). Interestingly, the number of microglia and their CD68 status seem to plateau at this stage, with no further increase in density beyond 12 months of age.

When the double homozygous TASTPM mice were examined, a very similar pattern was observed for densities of both total microglia and CD68 positive microglia (Fig 2d). As expected both plaque load and microglial response was greater in the homozygous mice.

Sections from single mutant mice were also stained for both IBA1 and CD68. The APP_Swe_ TAS mice, which develop plaques from around 16 months (Fig 1f), had a higher total and CD68 positive microglia at both 18 and 27 months of age (main effects of genotype for both Iba1 and CD68 in separate two-way ANOVAs; p<0.05). In contrast, the PSEN1_M146V_ TPM mice, which do not develop plaques at any age, showed no difference from wild type mice in their microglial phenotype at 4 months of age for either total or CD68 positive microglia (Fig 2g, two-tailed t-test, p>0.5).

Very similar changes in total microglial and CD68 positive microglial densities were found in dentate gyrus, CA3 and subiculum (data not shown).

### Synaptic currents in CA1 neurones

#### Spontaneous excitatory activity

When spontaneous excitatory currents are recorded from CA1 pyramidal neurones in the presence of a GABA_A_ receptor antagonist (gabazine, 6 μM), the substantial loss of spontaneous activity we have previously reported up to 4 months of age^[16]^ is largely maintained through to 18 months (Fig 3a). At 8 months there was almost a complete loss of action potential-dependent spontaneous activity in the transgenic mice (Fig 3aii), with the frequency almost exactly the same as that of miniature currents (Fig 3aiii) observed in the presence of 1 μM tetrodotoxin. This result was almost identical to what we have previously reported at 4 months. By 12-18 months, a decrement in frequency of miniature EPSCs had also developed but the action potential mediated spontaneous excitatory current frequency, although significantly lower in transgenic mice than wild type mice, were at a higher frequency than the miniature currents in the same genotype, implying that some spontaneous action potential-mediated activity was occurring in the transgenic mice at these older ages (Fig 3a). There were no differences between genotypes of mEPSC amplitudes or decay time constants (data not shown).

**Figure 3.**
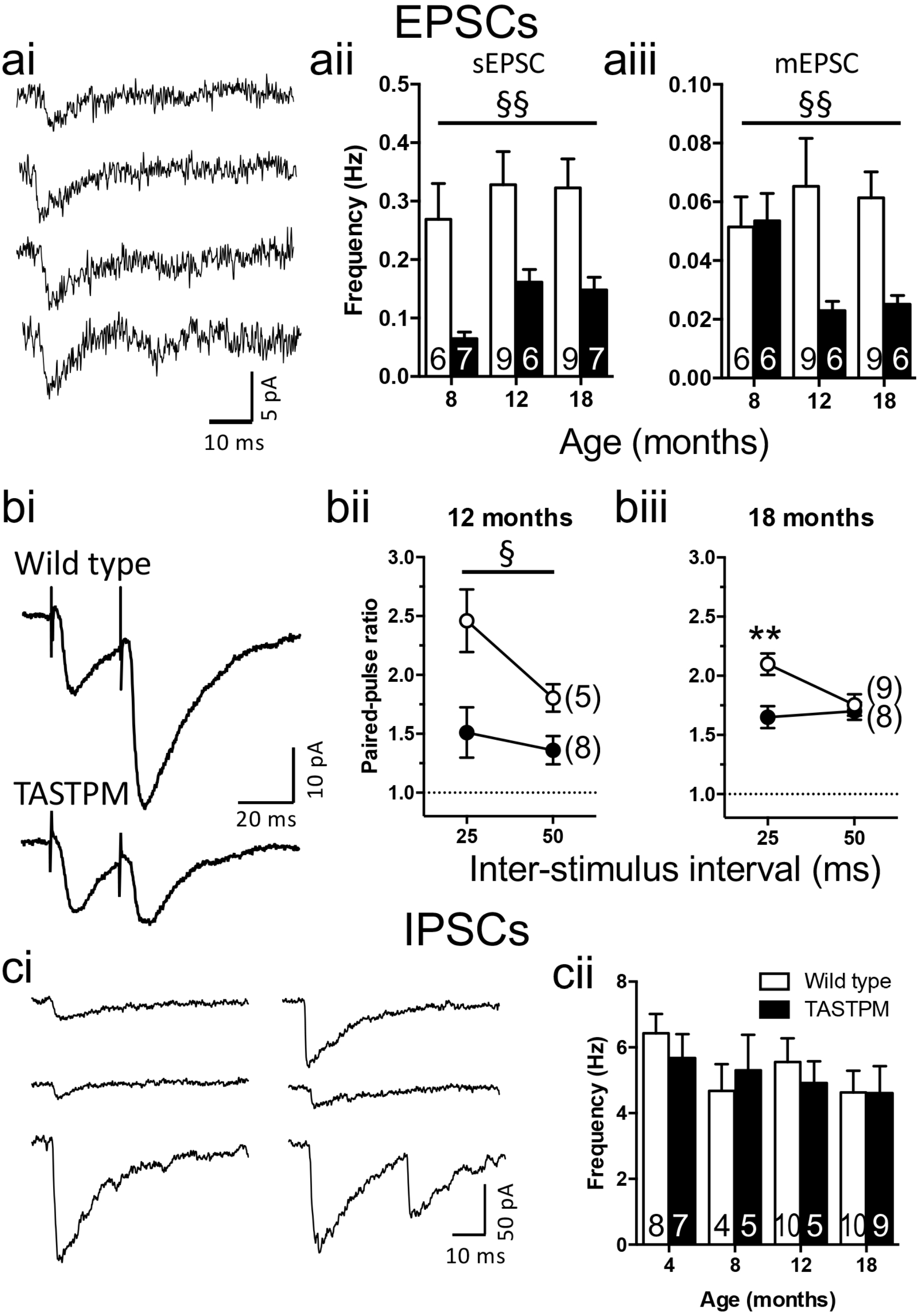
Excitatory and inhibitory synaptic transmission in TASTPM mice. ai) Example spontaneous and miniature EPSCs recorded from 18-month-old TASTPM. aii) Frequency of spontaneous EPSCs. Main effect of genotype by 2-way ANOVA (p=0.0011). aiii) Frequency of miniature EPSCs. Main effect of genotype by 2-way ANOVA (p < 0.01) bi) Example responses evoked by paired stimuli. bii) Paired-pulse ratios from 12-month-old mice. Main-effect of genotype by 2-way ANOVA (p=0.02). (biii) Paired-pulse ratios from 18-month-old mice. Genotype by inter-stimulus interval interaction (p<0.05); Sidak post-hoc test is indicated (p=0.002). ci) Example continuous recordings of spontaneous IPSCs. cii) Spontaneous IPSC frequencies. Sample sizes are indicated by the numbers within bars or within parentheses. * p<0.05; ** p<0.01. All data are represented by mean±SEM.

When excitatory activity was evoked by stimulating the Schaffer collaterals in slices from older animals, the previously reported increase in release probability at 4 months, as assessed by a lower paired-pulse facilitation and confirmed by a reduction in failure to release glutamate in response to minimal stimulation^[16]^, was maintained. Thus, at 12 months of age, paired-pulse ratio was lower in TASTPM than wild type mice at both 25 and 50 ms inter-stimulus intervals (n=5-8, two-way ANOVA, main effect of genotype p<0·02), while at 18 months of age, the lower paired-pulse ratio was only evident at 25 ms, resulting in a genotype x interval interaction (p<0·05) and a highly significant Sidak post hoc pairwise comparison between genotype at 25 ms (p<0·005).

#### Inhibitory activity is unchanged

Although we have concentrated our detailed analysis on glutamatergic activity, we were able to gain an overview of inhibitory activity from the spontaneous activity recorded in the absence of antagonists by utilising CsCl as the internal patch pipette solution. Hence, in the experiments examining pharmacologically isolated EPSCs above, initially spontaneous currents were recorded in the absence of receptor antagonists. When the GABA_A_ receptor antagonist was included in the perfusion solution, the frequency of currents was decreased to about 5-10% in wild type slices. Consequently, GABA_A_ receptor-mediated activity contributes about 90% of the frequency of the initial recording of mixed spontaneous activity. Note that, if the recording were made in the presence of a glutamatergic antagonist, any currents that were due to spontaneous glutamatergic activity mediating feed-forward or feed-back inhibition would be lost. Hence, the assessment of the drug-free frequency is a better assessment of the contribution of inhibition with the understanding that this will overestimate frequency by about 10%. Under these conditions we find that there is no significant change in inhibitory synaptic activity in TASTPM mice at any age from 4 to 18 months (Fig 3c). Note that, considering the very substantial change in spontaneous glutamatergic activity, this implies that very little of the GABAergic activity is dependent on glutamatergic input.

#### Synaptic plasticity is altered in TASTPM mice

Considering the changes in excitatory transmission it seemed likely that synaptic plasticity, in particular, long-term potentiation (LTP), the best cellular model we have for the laying down and retrieval of memory^[20]^, could be altered. To assess effects on LTP, field potentials were recorded from CA3-CA1 synapses in acute brain slices prepared from TASTPM mice at ages preceding detectable Αβ plaques (2 months) through to ages with a heavy plaque load (12-18 months).

Field input-output relationship was not significantly changed at any of the ages tested (Fig 4a). Moreover, unlike our previously reported result using patch clamp recording in the presence of a GABA_A_ receptor antagonist^[16]^, in field recordings with the inhibitory network intact, paired-pulse ratios were not significantly changed between genotypes at any of the ages tested (Fig 4b). This suggests that the observed changes in paired-pulse ratio were masked when GABAergic activity was not blocked, presumably, at least in part, by feed-forward inhibition decreasing release on the second stimulus.

**Figure 4.**
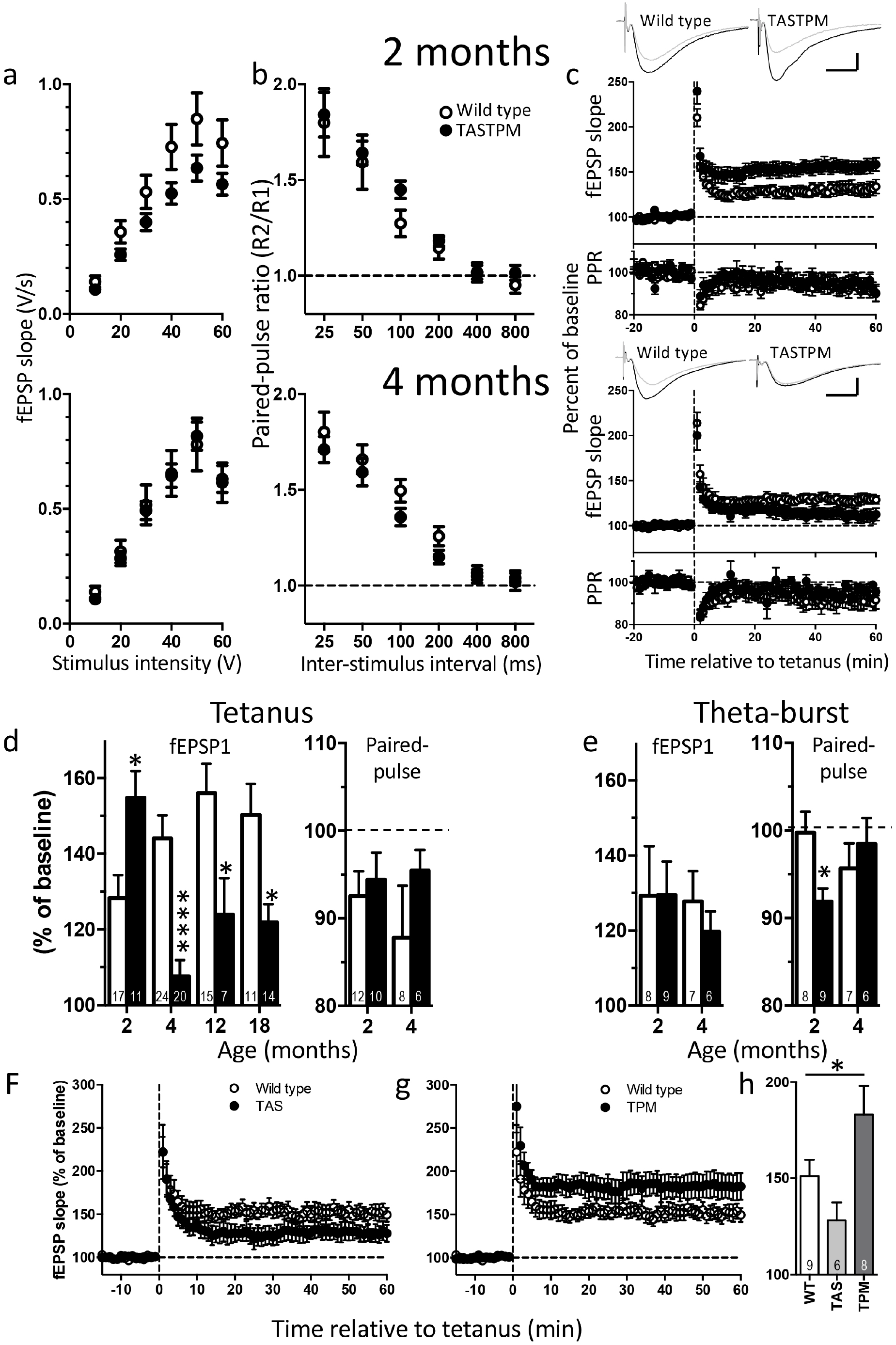
Field EPSP recordings of synaptic plasticity in TASTPM mice. a-c) 2 months upper row; 4 months lower row. a) Input-output relationship, b) Paired-pulse ratio profile and c) Time course of tetanus-induced LTP: fEPSP (upper panels) and paired-pulse ratio (PPR, lower panels) after induction as a percentage of pre-induction baseline. Traces are representative fEPSPs from baseline (grey) and the 51-60 minutes after tetanus (black). Scale bars: 5 ms × 0.5 mV. d) Magnitude of LTP (*left*) induced by tetanus over 2-18 months of age (number of mice as indicated in each column). Two-way ANOVA revealed significant age x genotype interaction (P<0.0001). Change in paired-pulse ratio (*right*) following LTP induction. Both wild type and TASTPM mice showed a significant decrease in PPR compared to baseline at both ages (paired t-tests within separate experiments; p < 0.05). No significant difference between genotypes (two-way ANOVA). e) Magnitude of LTP induced by theta-burst stimulation (*left,* no significant difference between genotypes) and corresponding change in paired-pulse ratio (*right*; two-way ANOVA age x genotype interaction p < 0.05). Sidak post hoc test: *P<0.05; ****P<0.0001. f) LTP induced by tetanic stimulation in APP_Swe_ TAS mice at 24 months of age. g) LTP induced by tetanic stimulation in PSEN1_M146V_ TPM mice at 24 months. Wild type animals are common to both panels f and g. h) Magnitude of LTP induced in 24 month old wild type, TAS and TPM mice. 24 month old recordings for all genotypes were interleaved; significance tested by one-way ANOVA, *p<0.05. All data are represented by mean±SEM.

At 2 months of age, LTP is altered in TASTPM mice but this is dependent on the induction protocol. When LTP was induced by tetanic stimulation in these young mice, the magnitude of LTP was greater in slices from TASTPM than from wild type mice. Paired-pulse ratios remained unaltered when the last ten minutes of the post-induction recordings was compared to the baseline in both genotypes (Fig 4c&d). This suggested that the change in amplitude seen in LTP was due to a postsynaptic change. In contrast, when LTP was induced using TBS, there was no difference between the wild type and transgenic mice (Fig 4e). Surprisingly, however, despite LTP remaining unchanged after theta burst stimulation, the paired-pulse ratio decreased significantly between the pre-induction baseline and the last 10 minutes of LTP in the TASTPM but not in the wild type mice. This suggests that the locus of expression of this form of LTP is likely to have a presynaptic component in transgenic mice at this age. Note that, considering an increase in release probability would be expected to enhance the postsynaptic response, this would suggest that the postsynaptic contribution to LTP could be decreased in the transgenic mice possibly with one locus of change compensating for the other.

At 4 months of age, the effects of the different stimulus protocols had changed. By this age, a clear deficit was seen in LTP induced by tetanic stimulation, which was the reverse change to that seen at 2 months. Moreover, similar effects were seen in two separate cohorts of 4-month-old mice (cohort 1 shown in Fig 4c; pooled data shown in Fig 4d.

TBS-induced LTP (Fig 4e) was, like at 2 months, of similar magnitude between genotypes but analysis of paired-pulse ratios, following LTP induction, indicated that LTP has a similar locus of expression in the two genotypes.

In addition, chemically-induced LTP was also examined in a separate set of slices, using transient application of the potassium channel blocker tetraethylammonium, applied after completion of a tetanus-induced LTP experiment. As expected, tetraethylammonium application resulted in an initial reduction in the field potential, followed by a long-lasting increase after wash-out. The magnitude of the tetraethylammonium-induced LTP was unchanged in slices of TASTPM compared to wild type animals (data not shown). This confirms that, despite deficits in tetanus-induced LTP, TASTPM are capable of expressing LTP under other induction protocols.

The deficit in tetanus-induced LTP was maintained at 12 and 18 months of age (Fig 4e).

#### Synaptic plasticity in TAS and TPM mice

The induction of LTP was also examined in 24 month old single mutant mice (Fig 4f-h). In age-matched wild type mice, LTP magnitude was similar to that observed at younger ages. In the APP_Swe_ TAS mice, the magnitude of LTP was smaller than wild type mice (Fig 4f). In contrast, in the PSEN1_M146V_ TPM mice, the magnitude of LTP was greater (Fig 4g). There was an overall significance between genotypes by one-way ANOVA (p<0.05; Fig 4h, Fisher LSD *post hoc* tests versus wild type p=0.1 for TAS and p=0.05 for TPM).

### Cognition and behaviour

#### TASTPM mice show rigidity in hippocampus-dependent learning

The hippocampus-dependent forced-alternation T-maze task was used to assess cognition in the TASTPM mice at three ages (Fig 5): 4 months, which corresponds to the first appearance of Αβ plaques (Fig 1) and where tetanus induced LTP was first impaired (Fig 4); 8 months, a moderate plaque load and 12 months, a heavy plaque load. Due to availability of animals, in this set of experiments, the 4-month-old group was run as three cohorts (no significant difference between the cohorts); while the 8- and 12-month-old group were a single cohort that underwent repeat training in a longitudinal study. It should be noted that at both 4 and 8 months of age, a subset of TASTPM mice did not complete the task in the required time and these mice were not included in the final results (1 of 16 mice at 4 months of age and 3 of 13 mice at 8 months of age).

**Figure 5.**
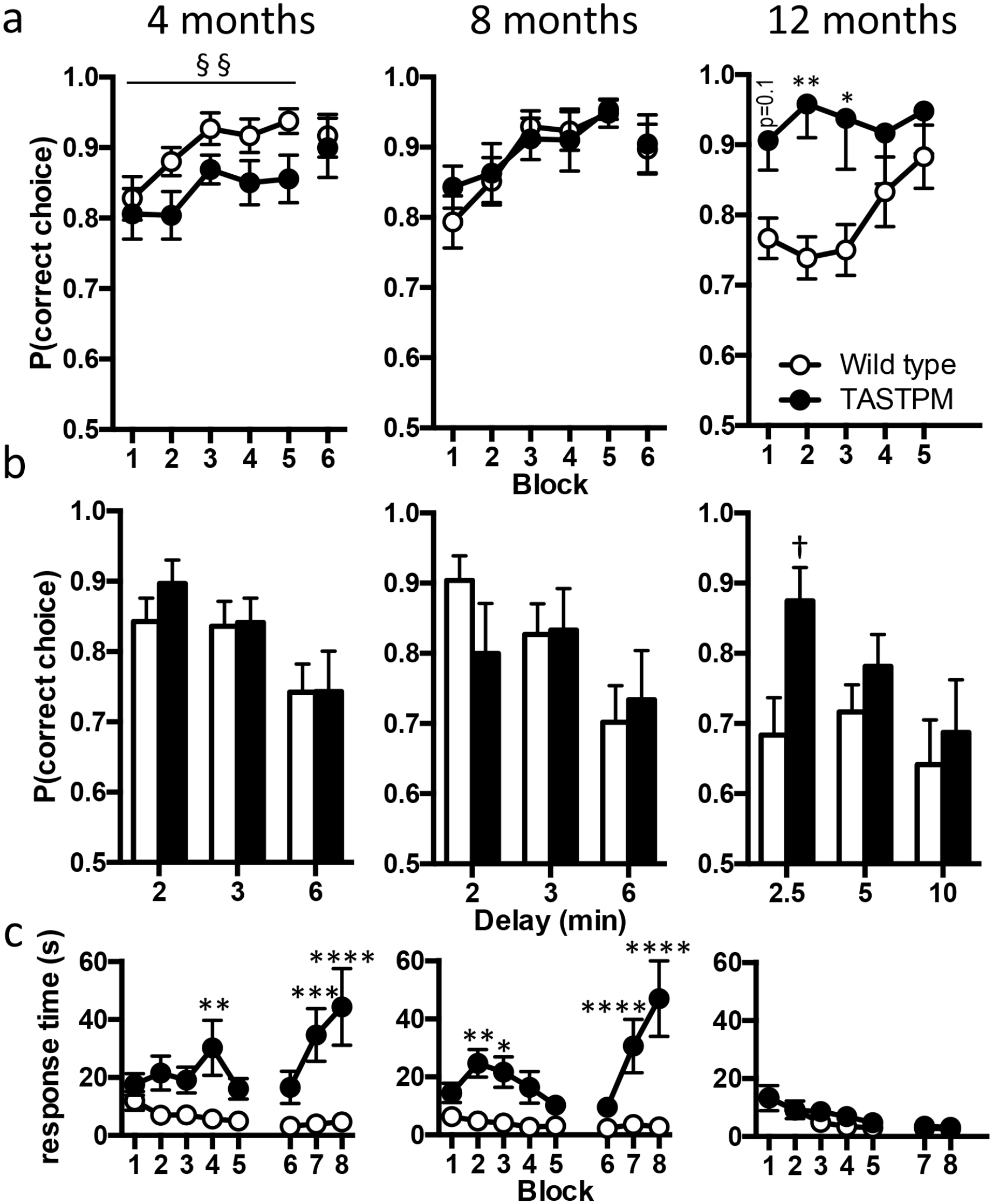
Hippocampus-dependent T-Maze learning in TASTPM mice. a-c) Ages as indicated above panel a. a) Probability of mice making a correct choice in the choice run at 4 months (n=15-16 animals completing the task), 8 months (n=10-13 animals completing the task) and 12 months of age (wild type n=15; TASTPM n=8). At 4 months of age there was a main effect of genotype in a two-way ANOVA (§§ p<0.01). Note the difference in starting performance at 12 months of age. These are the same mice tested at 8 months. As expected wild type mice started at around 75% correct choices and improved with training having, extinguished their previous training. TASTPM mice unexpectedly started training making correct choices 90% of the time having retained the previous improved level. Two-way ANOVA interaction between training block and genotype (p<0.03); Sidak post hoc analyses are indicated by *P<0.05, **P<0.01. b) The introduction of delays between sample and choice runs revealed no further differences between genotypes at 4 or 8 months of age, while at 12 months, a significant effect of genotype was observed with TASTPM mice retaining performance despite a 2.5 minute delay before the choice run († p<0.05). c) Response time for choice run at 4 -12 months of age, Sidak post hoc analyses are indicated by *P<0.05, **P<0.01, ***P<0.001, ****P<0.0001. All data are represented by mean±SEM.

At 4 months of age, both genotypes started training at similar levels of performance and improved over the training period (Fig 5a, two-way ANOVA, main effect of 5 block training period, p=0·01). However, TASTPM performed significantly worse than wild type mice overall (main effect of genotype, p<0·003), and there was no significant interaction between training block and genotype. Interestingly in Block 6, after 2 days without training, the two groups came together with almost identical performance at 90% correct choices (Fig 5a). When extra delays (2-6 minutes) were then added between the sample and choice runs a slight decrement in performance occurred, as expected, but there was no difference between genotypes (Fig 5b). In terms of the behaviour within the trials it was notable that on both the sample and choice runs the TASTPM mice showed considerably longer response times. In the case of the choice run, this may have influenced the result, especially the apparent impairment during the training period, as the time that memory needed to be retained was more than doubled (Fig 5c).

At 8 months of age, TASTPM performed similarly to wild type mice, improving over the training period (two-way ANOVA, main effect of training period (p<0·0001). When the same mice that were run at 8 months were retested at 12 months of age, wild types had, as expected, returned to the baseline response of approximately 75% correct and again improved from ~75% to ~90% during the training period. Unexpectedly however, TASTPM appeared to have retained their previous training, starting the training at ~90% and showed no further improvement over the training period. This was reflected in a significant interaction between training period and genotype in a two-way ANOVA (p<0·03, Fig 5a).

Animals were again challenged further by the introduction of a delay between sample and choice runs (Fig 5c). While at 8 months there was no difference between genotypes, at 12 months of age although TASTPM mice showed a similar decrement with the longest delays their performance with a 2·5 minute delay between sample and choice runs was surprisingly better than their wild type counterparts (two-tailed t-test, p<0·05).

Another interesting difference between the 12-month mice and the younger groups was that the response time for the choice run for TASTPM mice at 12 months was rapid and no different from the wild type mice (Fig 5c). (Note, at this age the mice also always completed the task in the required time.) In contrast, as described for 4 months above, the 8-month-old TASTPM mice showed a considerable delay to respond. However, despite this delay again increasing the time that memory needed to be retained, the 8 months mice showed no decrement in correct choices.

These failures and delays to respond may be interesting in terms of early behavioural changes and may also suggest different levels of anxiety.

#### TASTPM mice show decreased activity and increased anxiety-related behaviour compared to wild type mice

In the light of the behavioural differences seen in the T-maze trials, locomotor activity and anxiety were assessed using a battery of tests. First, mice were placed in an open field (Fig 6a). TASTPM mice had shorter total path lengths than their wild type counterparts at all ages (Fig. 6ai; two-way ANOVA, main effect of genotype, p<0·0001; no interaction between age and genotype). Furthermore, TASTPM mice spent more time in the peripheral than central area (Fig. 6aii; two-way ANOVA, main effect of genotype, p<0·0005; main effect of age, p<0·0001).

**Figure 6.**
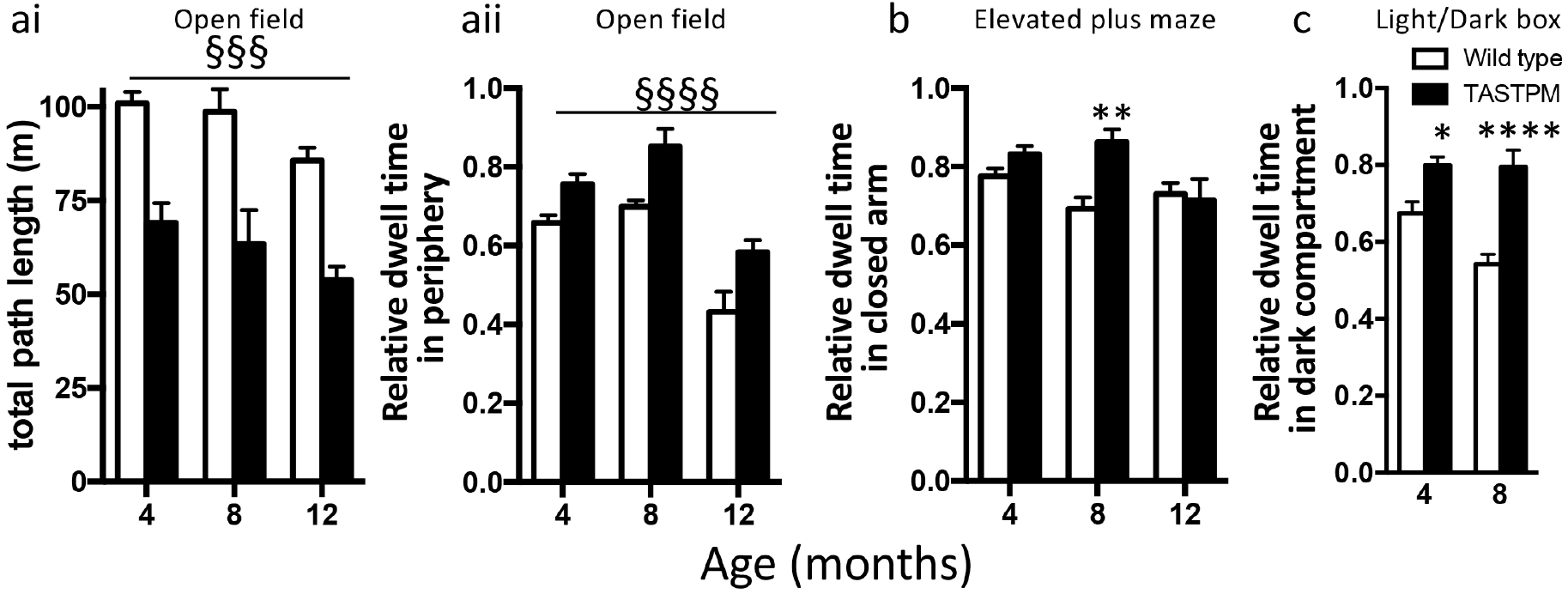
Open field and tests of anxiety. a) Open field test. ai) Total path length run over 30 mins. Main effect of genotype by two-way ANOVA (§§§ p<0.001). aii) Time spent in peripheral area. Main effect of genotype by two-way ANOVA (§§§§ p<0.0001). b) Elevated plus maze. Relative time in closed arm. c) Light/dark box. Relative time spent in dark compartment (12 months not tested.) In b and c, Sidak post hoc tests are indicated (two-way ANOVA genotype x age interaction P<0.05; * p<0.05; ***p<0.001; **** p<0.0001). All data are represented by mean±SEM.

Secondly, at 8 months of age, TASTPM mice spend more time in the closed than open arms of an elevated plus maze (two-way ANOVA interaction between age and genotype p<0·05); at 4 and 12 months of age this difference was not evident (Fig. 6b).

Finally, at 4 and 8 months of age, TASTPM mice spent more time in the closed compartment of a light/dark box (12 months not tested). There was a significant interaction between age and genotype (Fig. 6c; p<0·05).

Overall, these data indicate an increased level of anxiety in TASTPM mice from early stages of plaque development especially at earlier ages.

## Discussion

### Pathology and microglia

Proliferation of microglia, particularly clustering around plaques, has been repeatedly reported in Alzheimer’s disease, both in the human condition and in mouse models.^[21-23]^ Moreover we have previously reported a close correlation between plaque load and a microglial coexpression module in TASTPM mice. This reveals not only changes in expression of the gene for the phagocytic protein CD68, tested with immunohistochemistry in the present study but also even greater changes in *Trem2* expression,^[10]^ a gene shown in humans to be important in Alzheimer’s disease in GWAS.^[24, 25]^ However, here we show increased density of microglia, even before plaques can be detected in this mouse model. This finding required immunohistochemistry and cell counts as the gene expression changes of such specific microglial genes would only be apparent in whole tissue analysis with very large changes, such as occurs in later stages. In the light of previous reports,^[21-23]^ proliferation seems the most likely explanation of the increased density, although other causes such as increased survival of microglia or migration into the hippocampus, are possible. The triggering of microglial proliferation may be due to soluble Αβ in the wider neuropil, as seem to be the case for the earliest synaptic changes.^[16]^ Alternatively, it is possible that initial seeds of plaques are already present at this early stage but are too small to detect, being below the detection threshold set here (<10 μm^2^).

In either case, the question arises as to how plaque development then continues and how this relates to increased microglial numbers and development of a reactive phagocytic phenotype. Once plaques are seeded at these early stages, the ongoing deposition may primarily contribute either to the growth of plaques or to the seeding of new plaques. It is also possible that large plaques are not the result of smaller plaques growing but rather due to the fusion of smaller plaques. Here we observe that the smallest plaques initially increase in number, almost doubling each month. This rate continues to increase, albeit not quite so rapidly, at least until about 14 months of age when the number of plaques plateaus. This suggests that small plaques are seeded throughout the ages tested and gradually grow to form the larger plaques. Presumably the rapid increase in the seeding of plaques during the early stages relates to the rapid increase in Aβ levels at this stage that we have previously reported.^[16]^ The plateauing of density of smaller plaques may also reflect them being gradually being engulfed as the larger plaques spread and thus they are no longer seen as separate entities. This is strongly suggested by the appearance of brighter spots of Aβ in the halo around some of the larger plaques as can be seen in Fig 1a from 10 months onwards.

As plaques form, we have previously shown a very strong correlation with microglial gene expression.^[10]^ This encompasses both proliferation and activation of microglia. While there are more microglia very early in these Αβ mice, their phagocytic phenotype appears later and seems only to be triggered once the larger plaques of >500 μm^2^ develop. However, it is important to note that a step up in the density of total and CD68 positive microglia also occur at this stage in the wild type mice. Although the number of microglia and their activation state remains higher in the TASTPM mice the similarity in the timing of this later stage in both genotypes suggests that this has a strong age-related component rather than being purely dependent on the presence of plaques. However, it is also interesting to note that in the homozygous TASTPM, which develop plaques earlier but plaque load plateaus at the same level as the hemizygous mice, the density of total microglia and CD68 positive microglia is higher than in hemizygotes. Furthermore, the absence of a microglial phenotype in the TPM mice, while the TAS mice show a similar phenotype excludes a direct effect of the mutant presenilin on microglial function.

As has been previously shown in TASTPM^[10]^ and other transgenic mice (for example, see reference 22), the microglia are strongly attracted to and cluster tightly around the plaques as they grow. The question arises as to what the microglia are actually doing in the Αβ mice. The assumption previously has been that they are removing or limiting expansion of plaques.^[22]^ Conversely, it has been repeatedly reported that removal of microglia by blocking CSF1R has no effect on plaque load,^[26-28]^ although a role for microglia in initial plaque deposition has been suggested,^[29]^ which may be related to the initial early rise in microglia numbers, even preceding detection of plaques. It is, however, clear in the TASTPM mice that, since plaques continue to grow in both size and number in parallel to rapid microglial proliferation, it does not seem that microglia are very effective at removing Aβ. Recently interesting evidence has been presented that Aβ stimulates microglia to phagocytose synapses in a complement dependent manner at early stages of plaque development in J20 (*APP*_Swe/Ind_) mice^[30]^ leading to the hypothesis that microglia are causing early synaptic damage. We put forward an alternative hypothesis that the microglia are rather removing spines and boutons damaged by the high levels of soluble Αβ in or near the plaques and that this may represent a protective function, avoiding further damage to the dendrites and axons affected and hence protecting network function. This is further supported by the observation that as plaques grow synaptic loss is concentrated near to the plaques and the loss decreases further from plaques.^[31, 32]^

### Synaptic transmission

At 8 months of age the synaptic changes observed are almost identical to those we have previously reported at 4 months, with no change in miniature currents but a complete loss of spontaneous activity and an increase in release probability of glutamate. Particularly at early stages, the plaque load in terms of the percentage tissue coverage is very low and so very few of the synapses recorded would be close to a plaque. Moreover, the observation that the change in paired-pulse ratio is similar despite a rising plaque load also suggests that the plaques themselves are unlikely to be causing this effect. Hence, this is presumably the result of very low levels of soluble Αβ throughout the tissue. Increased release probability has been reported to be the physiological effect of Αβ release in wild type rats.^[5]^ As the plaque load increases and larger plaques start to appear, additional changes become evident. At 12 months, although the evoked release probability is still similarly affected, being higher in the transgenic mice than the wild type mice, there is also a reduction in the frequency of miniature EPSCs. The simplest interpretations of loss of miniature frequency is either a loss of functional synapses or a decreased release probability (see reference 33 for review). As we have demonstrated that release probability is increased, this suggests that there is a loss synapses is the likely explanation. In and around the plaques soluble and insoluble Αβ will be in equilibrium and with increased plaque coverage with age, at least some of the synapses will be directly affected by the high local concentration of Αβ in the close vicinity of a plaque which could lead to this loss. Note that unlike evoked currents, miniature synaptic currents can originate from any synapse in the tissue and hence some will be physically near plaques. The increasing effect as the plaque load rises, supports this hypothesis. This is again consistent with the observation that synaptic loss occurs with increased plaque load but that the loss is inversely proportional to distance from a plaque.^[31]^

It is notable that the decrease in frequency of miniature EPSCs observed at 12 months coincides with the increase in microglial number and activation but that it does not decrease any further by 18 months. This is consistent with the plaque load and indeed the microglial density largely plateauing by 14 months. The very effective microglial proliferation and subsequent phagocytic phenotype, associated with strong upregulation of microglial genes such as *Trem2* that we have previously reported at this age,^[10]^ may be one of the reasons why the progression of Alzheimer’s pathology to neurofibrillary tangles and neurodegeneration does not go further in these mice.

### Synaptic plasticity

The biphasic change in LTP induced by a mild tetanic stimulus at the early stages of pathology in Αβ mice is of interest. At 2 months of age, prior to detection of plaques, when LTP is enhanced in the Αβ mice, Aβ is certainly present, albeit at low levels.^[10, 16]^ By 4 months of age, when the first plaques are evident, Aβ levels are at least 10-fold higher than at 2 months. At this stage the opposite effect is observed with magnitude of tetanus-induced LTP much reduced compared to wild types. This LTP deficit was maintained both at 12 and 18 months of age. The biphasic pattern of change may reflect the increasing glutamate release probability which we have previously shown is greater at 4 months than at 2 months of age. If glutamate were more readily released during conditioning, the initial outcome may be a greater calcium influx through NMDA receptors and thus a larger magnitude LTP. However, as levels rise by 4 months and release probability increases further, depletion of the readily-releasable pool of synaptic vesicles during the first pulses of the conditioning train may result in a failure to release transmitter as the train continues, thus impairing LTP. Alternatively, the higher levels of Aβ may have other effects, interacting with NMDA receptors directly and impeding LTP (reviewed in reference 34). Moreover, Aβ is itself proposed to be released in an activity-dependent manner^[35-37]^ and hence, as its levels rise, resulting in increased release probability, this could lead to a positive feedback.

In concert with (or alternatively to) the direct physiological effects of low levels of Aβ, the early increase in LTP magnitude we report could be an interaction with the mutant presenilin 1. Expression of both normal and mutant presenilin 1 has direct effects on synaptic transmission and plasticity. For example, the A246E mutation leads to increased LTP magnitude at young ages,^[38, 39]^ while the L286V mutation increases LTP magnitude at young ages but decreases at older (14 month) ages.^[40]^ Here we show that the M146V mutation increases the magnitude of LTP at 24 months of age, which could be the underlying mechanism of the initial increased LTP observed in the TASTPM, until the later Aβ effects become dominant, decrementing the LTP.

Interestingly when theta burst stimulation was used for induction, during which the readily-releasable pool would have a chance to replenish, there was no significant deficit in LTP at 4 months. However, under these conditions at 2 months, we see a decrease in paired-pulse ratio following induction of LTP that remains throughout the experiment, indicating an increase in release probability. As outline above, the physiological effect of Aβ is to increase release probability, and so this suggests that the theta rhythm itself may be causing local release of Αβ at the stimulated synapses. Despite this apparent presynaptic increase in transmission, the magnitude of LTP remains unaltered, raising the possibility of a postsynaptic decrease balancing this effect. However, when TEA was applied to slices, resulting in a wide-spread depolarisation and thus release of all neurotransmitter vesicles (excitatory, inhibitory and modulatory), LTP was induced similarly to wild types confirming that some forms of long-term plasticity remain intact. In our hands, the locus of expression of LTP, induced by tetanus, tends to have a presynaptic component in both genotypes, indicated by a change in paired-pulse profile following induction.

### Behaviour

In this study, TASTPM mice showed no measurable cognitive deficits at any age, despite a substantial plaque load.

There were, however, clear differences in motor activity and anxiety related behaviours. The TASTPM mice were slower to perform the T-maze task than the wild type mice or failed to do so altogether. Furthermore, in other tests, decreased activity and increased anxiety were evident. Although not specifically measured, the TASTPM mice are also more aggressive than wild type mice.^[41]^

The improved retention of learning in TASTPM mice, requiring no repeat training when the same mice were tested twice at 8 and 12 months, may appear counterintuitive. However, similar outcomes in hippocampal learning have been reported in TASTPM mice at around 4 months, using a contextual fear conditioning regime. While mice were unimpaired in learning the task in these previous reports, they failed to extinguish their learnt fear behaviour when conditioning was reversed.^[13, 14, 42]^ In the present study, the difference was additionally noted, with the TASTPM mice also making more correct choices than the wild type mice when a delay between runs was introduced.

The subtlety of the cognitive changes reported here in the TASTPM mouse, even at stages where a heavy plaque load is present, is consistent with previous findings.^[13-15, 42]^ Even when other Αβ models are considered, cognitive deficits are limited and, where present, it is unclear how they truly translate to the human disease (for reviews, see references 43, 44). Moreover, this is consistent with the human condition, where, at the time of diagnosis, when cognitive deficits are first being detected, there is already a substantial reduction in hippocampal volume of up to 20%,^[45]^ whereas there is no apparent neuronal loss in TASTPM mice. Furthermore, in humans, cognitive decline correlates well with Tau PET markers and less with Aβ load.^[46]^ Given that Αβ mouse models are not a complete model of AD, i.e. lacking Tau tangle formation and neurodegeneration, which more faithfully correlate with cognitive decline, they should be considered as models of the preclinical disease, when Αβ is first deposited.

Here we bring together the progression of pathology, microglia, synaptic transmission and other functional changes in a preclinical mouse model of AD. In a following paper we compare these results in mice with only plaques and no neurofibrillary tangles to a transgenic mouse model of later stage dementia in which neurofibrillary tangles occur without plaques.^[47]^

From the present study we conclude that the earliest effects of Αβ on synaptic transmission are probably due to very low levels of soluble Αβ in the volume of the tissue, rather than being directly related to plaques. At this early stage plaques are extremely small and sparse so that they would impinge directly on very few synapses. However, while wide-spread soluble Αβ may also influence early microglial changes these may well be influenced by the earliest seeding of plaques. Later changes most probably relate to the increasing numbers of larger plaques with the cloud of high-level soluble Αβ surrounding them impinging on a, still limited, but much greater area. The direct effects of this localised high concentration of soluble Αβ include loss of active synapses and activation of microglia, likely involved in clearing damaged synapses near or within plaques via phagocytosis and thus preventing wider damage to the affected axons and dendrites. While seeding of new plaques seems to continue throughout the development of plaque pathology, it is only once large plaques are present that these latter changes occur. Moreover, the lack of cognitive deficits is similar to the human situation prior to the build-up of neurofibrillary tangles and neurodegeneration and thus suggests that TASTPM represent a good model of the prodromal phase of the disease.

## Data sharing

The datasets used and/or analysed during the current study are available from the corresponding author on reasonable request.

## Acknowledgements

The authors would like to thank Rivka Steinberg and Stuart Martin for providing genotyping services; Ken Smith, Roshni Desai and Tammaryn Lashley for histological advice; and the Maria Fitzgerald/Steve Hunt laboratories for use of equipment.

## Authors’ contributions

EM performed the histological staining and counting of microglia and provided feedback on draft manuscript. TB carried out behavioural experiments and analysis and comments on draft manuscript. WL performed the LTP experiments at 12 and 18 months of age and comments on draft manuscript. TH performed LTP and behavioural experiments at 4 months of age. PH carried out the chemical LTP experiments and comments on draft manuscript. OTJ performed patch clamp experiments for IPSCs at 4 months of age. KY performed the LTP experiments at 24 months. LM provided training and supervision of all behavioural experiments. FM performed LTP experiments at 2 and 4 months of age. MANS performed behavioural experiments at 4 and 8 months of age. GB carried out histological staining and quantification of plaques. RW performed the microglial histology and counting for the TPM mice. JCR developed the mouse model and comments on the draft manuscript. FC designed and supervised the behavioural experiments and provided comments on the draft manuscript. DAS supervised the histological experiments and commented on draft manuscript. DMC supervised all experiments, performed electrophysiological recordings, wrote first draft of the manuscript, performed all statistical analyses, prepared figures, edited and finalised manuscript. FAE obtained funding, designed experiments, edited and finalised manuscript. All authors read and approved the final manuscript.

## References

1. Cuyvers E, Sleegers K. Genetic variations underlying Alzheimer’s disease: evidence from genome-wide association studies and beyond. Lancet Neurol 2016;15: 857-68. DOI:10.1016/S1474-4422(16)00127-7.

2. Hardy J, Allsop D. Amyloid deposition as the central event in the aetiology of Alzheimer’s disease. Trends in Pharmacological Sciences 1991;12: 383-8.

3. Blennow K, Mattsson N, Scholl M, Hansson O, Zetterberg H. Amyloid biomarkers in Alzheimer’s disease. Trends Pharmacol Sci 2015;36: 297-309. DOI:10.1016/j.tips.2015.03.002.

4. Lewczuk P, Matzen A, Blennow K, et al. Cerebrospinal Fluid Abeta42/40 Corresponds Better than Abeta42 to Amyloid PET in Alzheimer’s Disease. J Alzheimers Dis 2017;55: 813-22. 10.3233/JAD-160722.

5. Abramov E, Dolev I, Fogel H, Ciccotosto GD, Ruff E, Slutsky I. Amyloid-beta as a positive endogenous regulator of release probability at hippocampal synapses. Nat Neurosci 2009;12: 1567-76. DOI:10.1038/nn.2433.

6. Dolev I, Fogel H, Milshtein H, et al. Spike bursts increase amyloid-beta 40/42 ratio by inducing a presenilin-1 conformational change. Nat Neurosci 2013;16: 587-95. DOI:10.1038/nn.3376.

7. Iaccarino HF, Singer AC, Martorell AJ, et al. Gamma frequency entrainment attenuates amyloid load and modifies microglia. Nature 2016;540: 230-5. DOI:10.1038/nature20587.

8. Verret L, Mann EO, Hang GB, et al. Inhibitory interneuron deficit links altered network activity and cognitive dysfunction in Alzheimer model. Cell 2012;149: 708-21. 10.1016/j.cell.2012.02.046.

9. Selkoe DJ, Hardy J. The amyloid hypothesis of Alzheimer’s disease at 25 years. EMBO Mol Med 2016;8: 595-608. DOI:10.15252/emmm.201606210.

10. Matarin M, Salih DA, Yasvoina M, et al. A genome-wide gene-expression analysis and database in transgenic mice during development of amyloid or tau pathology. Cell Rep 2015;10: 633-44. DOI:10.1016/j.celrep.2014.12.041.

11. Howlett DR, Bowler K, Soden PE, et al. Abeta deposition and related pathology in an APP x PS1 transgenic mouse model of Alzheimer’s disease. Histol Histopathol 2008;23: 67-76. DOI:10.14670/HH-23.67.

12. He Z, Guo JL, McBride JD, et al. Amyloid-beta plaques enhance Alzheimer’s brain tau-seeded pathologies by facilitating neuritic plaque tau aggregation. Nat Med 2018;24: 29-38. DOI:10.1038/nm.4443.

13. Rattray I, Scullion GA, Soulby A, Kendall DA, Pardon MC. The occurrence of a deficit in contextual fear extinction in adult amyloid-over-expressing TASTPM mice is independent of the strength of conditioning but can be prevented by mild novel cage stress. Behav Brain Res 2009;200: 83-90. DOI:10.1016/j.bbr.2008.12.037.

14. Rattray I, Pitiot A, Lowe J, et al. Novel cage stress alters remote contextual fear extinction and regional T2 magnetic resonance relaxation times in TASTPM mice overexpressing amyloid. J Alzheimers Dis 2010;20: 1049-68. DOI:10.3233/JAD-2010-091354.

15. Howlett DR, Richardson JC, Austin A, et al. Cognitive correlates of Abeta deposition in male and female mice bearing amyloid precursor protein and presenilin-1 mutant transgenes. Brain Res 2004;1017: 130-6. DOI:10.1016/j.brainres.2004.05.029.

16. Cummings DM, Liu W, Portelius E, et al. First effects of rising amyloid-beta in transgenic mouse brain: synaptic transmission and gene expression. Brain : a journal of neurology 2015;138: 1992-2004. DOI:10.1093/brain/awv127.

17. Richardson JC, Kendal CE, Anderson R, et al. Ultrastructural and behavioural changes precede amyloid deposition in a transgenic model of Alzheimer’s disease. Neuroscience 2003;122: 213-28. 10.1016/S0306-4522(03)00389-0.

18. Cacucci F, Yi M, Wills TJ, Chapman P, O’Keefe J. Place cell firing correlates with memory deficits and amyloid plaque burden in Tg2576 Alzheimer mouse model. Proc Natl Acad Sci U S A 2008;105: 7863-8. DOI:10.1073/pnas.0802908105.

19. Packard MG, Introini-Collison I, McGaugh JL. Stria terminalis lesions attenuate memory enhancement produced by intracaudate nucleus injections of oxotremorine. Neurobiol Learn Mem 1996;65: 278-82. 10.1006/nlme.1996.0033.

20. Bliss TVP, Collingridge GL, Morris RG. Synaptic plasticity in health and disease: introduction and overview. Philos Trans R Soc Lond B Biol Sci 2014;369: 20130129. DOI:10.1098/rstb.2013.0129.

21. Nahum-Levy R, Lipinski D, Shavit S, Benveniste M. Desensitization of NMDA receptor channels is modulated by glutamate agonists. Biophysical Journal 2001;80: 2152-66.

22. Condello C, Yuan P, Schain A, Grutzendler J. Microglia constitute a barrier that prevents neurotoxic protofibrillar Abeta42 hotspots around plaques. Nat Commun 2015;6: 6176. DOI:10.1038/ncomms7176.

23. Hong S, Dissing-Olesen L, Stevens B. New insights on the role of microglia in synaptic pruning in health and disease. Curr Opin Neurobiol 2016;36: 128-34. DOI:10.1016/j.conb.2015.12.004.

24. Guerreiro R, Wojtas A, Bras J, et al. TREM2 variants in Alzheimer’s disease. N Engl J Med 2013;368: 117-27. 10.1056/NEJMoa1211851.

25. Jonsson T, Stefansson H, Steinberg S, et al. Variant of TREM2 associated with the risk of Alzheimer’s disease. N Engl J Med 2013;368: 107-16. 10.1056/NEJMoa1211103.

26. Dagher NN, Najafi AR, Kayala KM, et al. Colony-stimulating factor 1 receptor inhibition prevents microglial plaque association and improves cognition in 3xTg-AD mice. J Neuroinflammation 2015;12: 139. DOI:10.1186/s12974-015-0366-9.

27. Spangenberg EE, Lee RJ, Najafi AR, et al. Eliminating microglia in Alzheimer’s mice prevents neuronal loss without modulating amyloid-beta pathology. Brain : a journal of neurology 2016;139: 1265-81. DOI:10.1093/brain/aww016.

28. Olmos-Alonso A, Schetters ST, Sri S, et al. Pharmacological targeting of CSF1R inhibits microglial proliferation and prevents the progression of Alzheimer’s-like pathology. Brain : a journal of neurology 2016;139: 891-907. 10.1093/brain/awv379.

29. Sosna J, Philipp S, Albay R, 3rd, et al. Early long-term administration of the CSF1R inhibitor PLX3397 ablates microglia and reduces accumulation of intraneuronal amyloid, neuritic plaque deposition and pre-fibrillar oligomers in 5XFAD mouse model of Alzheimer’s disease. Mol Neurodegener 2018;13: 11. DOI:10.1186/s13024-018-0244-x.

30. Hong S, Beja-Glasser VF, Nfonoyim BM, et al. Complement and microglia mediate early synapse loss in Alzheimer mouse models. Science 2016;352: 712-6. DOI:10.1126/science.aad8373.

31. Spires TL, Meyer-Luehmann M, Stern EA, et al. Dendritic spine abnormalities in amyloid precursor protein transgenic mice demonstrated by gene transfer and intravital multiphoton microscopy. J Neurosci 2005;25: 7278-87. DOI:10.1523/JNEUROSCI.1879-05.2005.

32. Kirkwood CM, Ciuchta J, Ikonomovic MD, et al. Dendritic spine density, morphology, and fibrillar actin content surrounding amyloid-beta plaques in a mouse model of amyloid-beta deposition. J Neuropathol Exp Neurol 2013;72: 791-800. DOI:10.1097/NEN.0b013e31829ecc89.

33. Edwards FA. Anatomy and electrophysiology of fast central synapses lead to a structural model for long-term potentiation. Physiological Reviews 1995;75: 759-87.

34. Zhang Y, Li P, Feng J, Wu M. Dysfunction of NMDA receptors in Alzheimer’s disease. Neurol Sci 2016;37: 1039-47. DOI:10.1007/s10072-016-2546-5.

35. Kamenetz F, Tomita T, Hsieh H, et al. APP processing and synaptic function. Neuron 2003;37: 925-37.

36. Cirrito JR, Kang JE, Lee J, et al. Endocytosis is required for synaptic activity-dependent release of amyloid-beta in vivo. Neuron 2008;58: 42-51. DOI:10.1016/j.neuron.2008.02.003.

37. Cirrito JR, Yamada KA, Finn MB, et al. Synaptic activity regulates interstitial fluid amyloid-beta levels in vivo. Neuron 2005;48: 913-22. DOI:10.1016/j.neuron.2005.10.028.

38. Parent A, Linden DJ, Sisodia SS, Borchelt DR. Synaptic transmission and hippocampal long-term potentiation in transgenic mice expressing FAD-linked presenilin 1. Neurobiol Dis 1999;6: 56-62. 10.1006/nbdi.1998.0207.

39. Dewachter I, Ris L, Croes S, et al. Modulation of synaptic plasticity and Tau phosphorylation by wild-type and mutant presenilin1. Neurobiology of Aging 2008;29: 639-52.

40. Auffret A, Gautheron V, Repici M, et al. Age-dependent impairment of spine morphology and synaptic plasticity in hippocampal CA1 neurons of a presenilin 1 transgenic mouse model of Alzheimer’s disease. JNS 2009;29: 10144-52.

41. Pugh PL, Richardson JC, Bate ST, Upton N, Sunter D. Non-cognitive behaviours in an APP/PS1 transgenic model of Alzheimer’s disease. Behav Brain Res 2007;178: 18-28. DOI:10.1016/j.bbr.2006.11.044.

42. Pardon MC, Sarmad S, Rattray I, et al. Repeated novel cage exposure-induced improvement of early Alzheimer’s-like cognitive and amyloid changes in TASTPM mice is unrelated to changes in brain endocannabinoids levels. Neurobiol Aging 2009;30: 1099-113. DOI:10.1016/j.neurobiolaging.2007.10.002.

43. Foley AM, Ammar ZM, Lee RH, Mitchell CS. Systematic review of the relationship between amyloid-beta levels and measures of transgenic mouse cognitive deficit in Alzheimer’s disease. J Alzheimers Dis 2015;44: 787-95. DOI:10.3233/JAD-142208.

44. Kobayashi DT, Chen KS. Behavioral phenotypes of amyloid-based genetically modified mouse models of Alzheimer’s disease. Genes Brain Behav 2005;4: 173-96. DOI:10.1111/j.1601-183X.2005.00124.x.

45. Ridha BH, Barnes J, Bartlett JW, et al. Tracking atrophy progression in familial Alzheimer’s disease: a serial MRI study. Lancet Neurol 2006;5: 828-34. DOI:10.1016/S1474-4422(06)70550-6.

46. Brier MR, Gordon B, Friedrichsen K, et al. Tau and Abeta imaging, CSF measures, and cognition in Alzheimer’s disease. Sci Transl Med 2016;8: 338ra66. DOI:10.1126/scitranslmed.aaf2362.

47. Joel Z, Izquierdo P, Liu W, et al. A TauP301L mouse model of dementia; development of pathology, synaptic transmission, microglial response and cognition throughout life. bioRxiv 2018;420398: DOI:10.1101/420398.

